# Frequency-dependent response of *Chromobacterium violaceum* to sonic stimulation, and altered gene expression associated with enhanced violacein production at 300 Hz

**DOI:** 10.1101/098186

**Authors:** Chinmayi Joshi, Pooja Patel, Abhishek Singh, Jinal Sukhadiya, Vidhi Shah, Vijay Kothari

## Abstract

*Chromobacterium violaceum* was subjected to sonic (100-2000 Hz) stimulation, and the effect on its cell yield and quorum sensing regulated pigment (violacein) production was investigated. Sound corresponding to the 300 Hz was found to promote (by 1.52 fold) violacein production the most, with only marginal impact on cell yield. Whole transcriptome analysis revealed that a total of 342 genes (i.e. 4.63% of whole genome) were significantly up-regulated in the sonic stimulated culture. Enhanced violacein production in the sound stimulated culture seems to have stemmed from enhanced expression of the genes involved in glucose metabolism through pentose phosphate pathway, resulting in increased availability of erythrose-4-phosphate, to be used for synthesis of tryptophan, the precursor for violacein synthesis. Multiple ribosomal subunit genes, enzyme coding genes, and those associated with secretion/transport were up-regulated owing to sonic stimulation. This study is a good demonstration of the ability of sound waves to alter bacterial metabolism.

## 1. Introduction

Though the interaction of microbial cells with audible range of sound has not been researched much, there are quite a few reports (Aggio et al., 2012; Cai et al., 2015; Sarvaiya and Kothari, 2015; Gu et al., 2016; Shah et al., 2016) describing microbial growth and metabolism getting affected by sound stimulation. Whether microbes respond to sonic stimulation in a frequency-dependent fashion, is an interesting problem to investigate. It is also possible that different microorganisms may respond differently to sound of same frequency. Microbes have been reported of being capable of producing sound and responding to it after sensing it (Matsuhashi et al., 1998). Sound has also been proposed as a possible signal for communication among microbes (Trushin, 2003; Reguera, 2011). Only limited knowledge is available regarding the influence of sonic waves on single cells and cellular metabolism. Audible sounds passing through the microbial growth medium in form of oscillating pressure waves can stimulate mechanosensory cells (Pickett et al., 2000).

Quorum sensing (QS) is the term used to describe the phenomenon whereby microbial cells communicate among themselves. This process is believed largely to be mediated through the chemical signals like acyl homoserine lactone (AHL), or small peptide signals. Many of the traits including pigment production among bacteria have been shown to be regulated by QS. *Chromobacterium violaceum* has been one of the most widely used model bacterium for study of QS in gram-negative bacteria. Production of the purple pigment violacein in this bacterium is under control of its QS machinery (Ghosh et al., 2014). Violacein is a bioactive molecule with potential commercial and therapeutic importance. Various research groups are seeking to improve the fermentative yields of violacein through genetic/ metabolic engineering and synthetic biology (Choi et al., 2015; Xu et al., 2016).

In the present study, we report our results regarding influence of audible sound (100-2000 Hz) on growth and QS-regulated pigment production of the bacterium *C. violaceum*.

## 2. Materials and Methods

### 2.1. Bacterial culture

*Chromobacterium violaceum* (MTCC 2656) was procured from the Microbial Type Culture Collection (MTCC), Chandigarh. This organism was grown in nutrient broth (HiMedia, Mumbai) supplemented with 1% v/v glycerol (CDH^®^ New Delhi). Incubation was made at 35° C.

### 2.2. Sound generation

Sound beep(s) of required frequency was generated using NCH^®^ tone generator. The sound file played during the experiment was prepared using Wave Pad Sound Editor Masters Edition v.5.5 in such a way that there is a time gap of one second between two consecutive beep sounds.

### 2.3. Sound stimulation

Inoculum of the test bacterium was prepared from its activated culture, in sterile normal saline. Optical density of this inoculum was adjusted to 0.08-0.10 at 625 nm (Agilent Technologies Cary 60 Uv-Vis). The test tubes (Borosil, 18 X 150 mm; 27 mL) containing 6 mL of growth medium (including 5% v/v inoculum) were put into a glass chamber (Actira, L: 250 x W: 150 x H: 250 mm). A speaker (Lenovo) was put in this glass chamber at the distance of 15 cm from the inoculated test tubes. Sound delivery from the speaker was provided throughout the period of incubation (48 h). This glass chamber was covered with a glass lid, and one layer of loose-fill shock absorber polystyrene, in such a way that the polystyrene layer gets placed below the glass lid. Silicone grease was applied on the periphery of the glass chamber coming in contact with the polystyrene material. This type of packaging was done to minimize any possible leakage of sound from inside of the chamber, and also to avoid any interference from external sound. Similar chamber was used to house the ‘control’ (i.e. not subjected to sound stimulation) group test tubes. One speaker was also placed in the glass chamber used for the control tubes at a distance of 15 cm, where no electricity was supplied and no sound was generated (Kothari et al., 2016). Intensity of sound, measured with a sound level meter (ACD machine control Ltd.) at a distance of 15 cm from the speaker at each of the test frequency has been listed in Table-1. Sound level in control chamber was found to be below the detection level (<40 dB) of the sound level meter. Schematic of the whole experimental set-up is shown in Figure 1.

**Figure 1.**
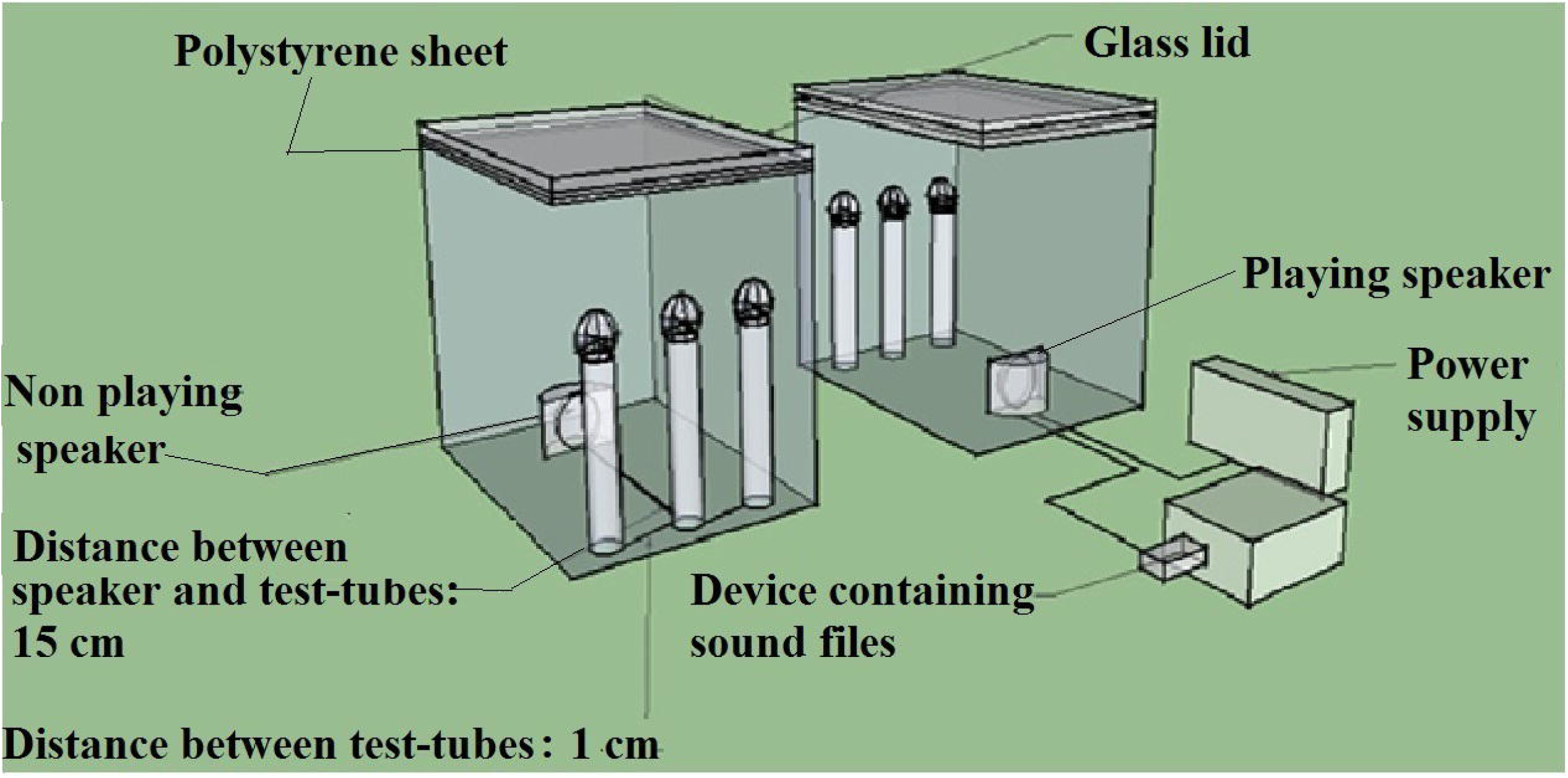
Schematic of the experimental set-up.

Intermittent mixing of the contents of the test tubes to minimize heterogeneity was achieved by vortexing the tubes at an interval of every 3 h using a cyclomixer. Whenever the tubes were taken out for cyclomixer, each time positions of tubes of a single chamber were inter-changed, and their direction with respect to speaker was changed by rotating them 180°. This was done to achieve a high probability of almost equal sound exposure to all the tubes.

### 2.4. Growth and pigment estimation

At the end of incubation, after quantifying the cell density at 764 nm (Joshi et al., 2016), the culture tubes were subjected to extraction of the pigment violacein (Choo et al, 2006). Briefly, 1 mL of the culture broth was centrifuged (REMI CPR-24 Plus) at 12,000 rpm (13,520 g) for 15 min at 25°C, and the resulting supernatant was discarded. The remaining cell pellet was resuspended into 1 mL of DMSO (Merck, Mumbai), and incubated at room temperature for 30 min, followed by centrifugation at 12,000 rpm for 15 min. The violacein extracted in supernatant was estimated by measuring OD at 585 nm. Violacein unit was calculated as OD_585_/OD_764_. This parameter was calculated to nullify the effect of change in cell density on pigment production.

### 2.5. Gene expression analysis

Before RNA isolation, taxonomic identity and purity of culture was confirmed by 16srRNA sequencing.

#### 2.5.1. Total RNA isolation, and its qualitative and quantitative analysis

Total RNA was isolated from the samples using Trizol (Invitrogen) method for RNA extraction.The quality of the total RNA was checked on 1% denatured agarose gel and quantified using Qubitfluorometer. The samples were treated with *MICROB Express* kit to deplete the bacterial ribosomal RNA and enrich the mRNA population to be used for whole transcriptome analysis (WTA) library preparation. The enriched mRNA was fragmented and the libraries were prepared using *illuminaTruSeq RNA Sample Preparation V2 kit*. The means of the library fragment size distributions were 725 and 734 bp for ‘control’ and ‘experimental’ samples respectively. The libraries were sequenced using 2x150 PE chemistry generating ~1.5-2.0 GB of data per sample.

#### 2.5.2. Illumina 2 x 150 PE library preparation, Cluster Generation and Sequencing

The samples were initially treated with *MICROBExpress* kit (Ambion/Life tech) to deplete the ribosomal RNA and enrich the bacterial mRNA population. Enriched bacterial mRNA was then reverse transcribed into first-strand cDNA using SuperScript II reverse transcriptase (Invitrogen) and random hexamer primer as per *illuminaTruSeq RNA Sample Preparation V2 kit* protocol. The single stranded cDNA was then converted to ds cDNA. ds cDNA samples were subjected to Covaris shearing followed by end-repair of overhangs resulting from shearing. The end–repaired fragments were A-tailed, adapter ligated and then enriched by limited number of PCR cycles. Library quantification was performed using DNA high sensitivity assay kit. The next generation sequencing for control and experimental samples was performed on the Illumina platform and approximately ~5-6 GB of data was been generated per sample. The high quality pair end reads of both the samples were assembled together using SOAP*denovo-Trans* (version-1.03). The final assembly contained 779 transcripts with an N50 scaffold length of 22,087 bp; the largest transcript assembled measured 119,734 bp. A total of 7,403 coding regions were identified using TransDecoder (version-2.0.1) program. The amplified libraries were analyzed in Bioanalyzer 2100 (Agilent Technologies) using HighSensitivity (HS) DNA chip as per manufacturer’s instructions.

After obtaining the Qubit concentration for the libraries and the mean peak size from Bioanalyser profile, library was loaded into Illumina platform for cluster generation and sequencing. Paired-end sequencing allows the template fragments to be sequenced in both the forward and reverse directions. The library molecules bind to complementary adapter oligos on paired-end flow cell. The adapters are designed to allow selective cleavage of the forward strands after re-synthesis of the reverse strand during sequencing. The copied reverse strand was used to sequence from the opposite end of the fragment.

### 2.6. Functional Annotation of Predicted Protein Coding Genes

Each of the predicted protein coding genes were annotated evaluating the homology by BLASTX search against NR (Non-redundant) database. To identify the biological pathways that are active in both (control and experimental) samples, the predicted ORFs were mapped to reference canonical pathways in KEGG using KEGG automatic annotation server (KAAS).The DESeq package was used to identify significantly differentially expressed genes between control and experimental samples.

### 2.7. Statistical analysis

All the experiments were performed in triplicate, and measurements are reported as mean ± standard deviation (SD). Statistical significance of the data was evaluated by applying t-test using Microsoft excel^®^. Data with *p*-value less than 0.05 was considered to be statistically significant.

## 3. Results and Discussion

### 3.1. Effect of different sonic frequencies on C. violaceum

Response of *C. violaceum* to sound stimulation at different frequencies in terms of growth and pigment production is summarized in Figure 2. Out of nine test frequencies, eight caused significant change in bacterial growth. In all these eight cases the effect on growth was stimulatory. Significant change in violacein unit (VU) was observed at seven of the test frequencies. Significant change in VU can be considered as an indication of true QS-modulatory effect of the sound treatment, largely independent of the alteration in growth. In five cases (i.e. 300 Hz, 400 Hz, 600 Hz, 700 Hz, and 2,000 Hz) magnitude of alteration in QS-regulated violacein production owing to sound stimulation, was much higher than that in cell density. This is to say that the QS-regulated trait (i.e. pigment production) got affected more than the bacterial growth. Among the five QS-modulatory sound frequencies, maximum (51.28%) positive effect on QS-regulated pigment production with least (4.54%) effect on growth was observed at 300 Hz. Looking at the notable effect of 300 Hz sonic stimulation on bacterial QS, we subjected the cells exposed to 300 Hz sound to transcriptome profiling, along with the ‘control’ cells, which received no sound stimulation. The QS-regulated trait i.e. violacein production was maximally affected at 300 Hz and 2,000 Hz; however effect of 2,000 Hz sound on VU being negative, sample treated with 300 Hz sound was selected for whole transcriptome analysis.

**Figure 2.**
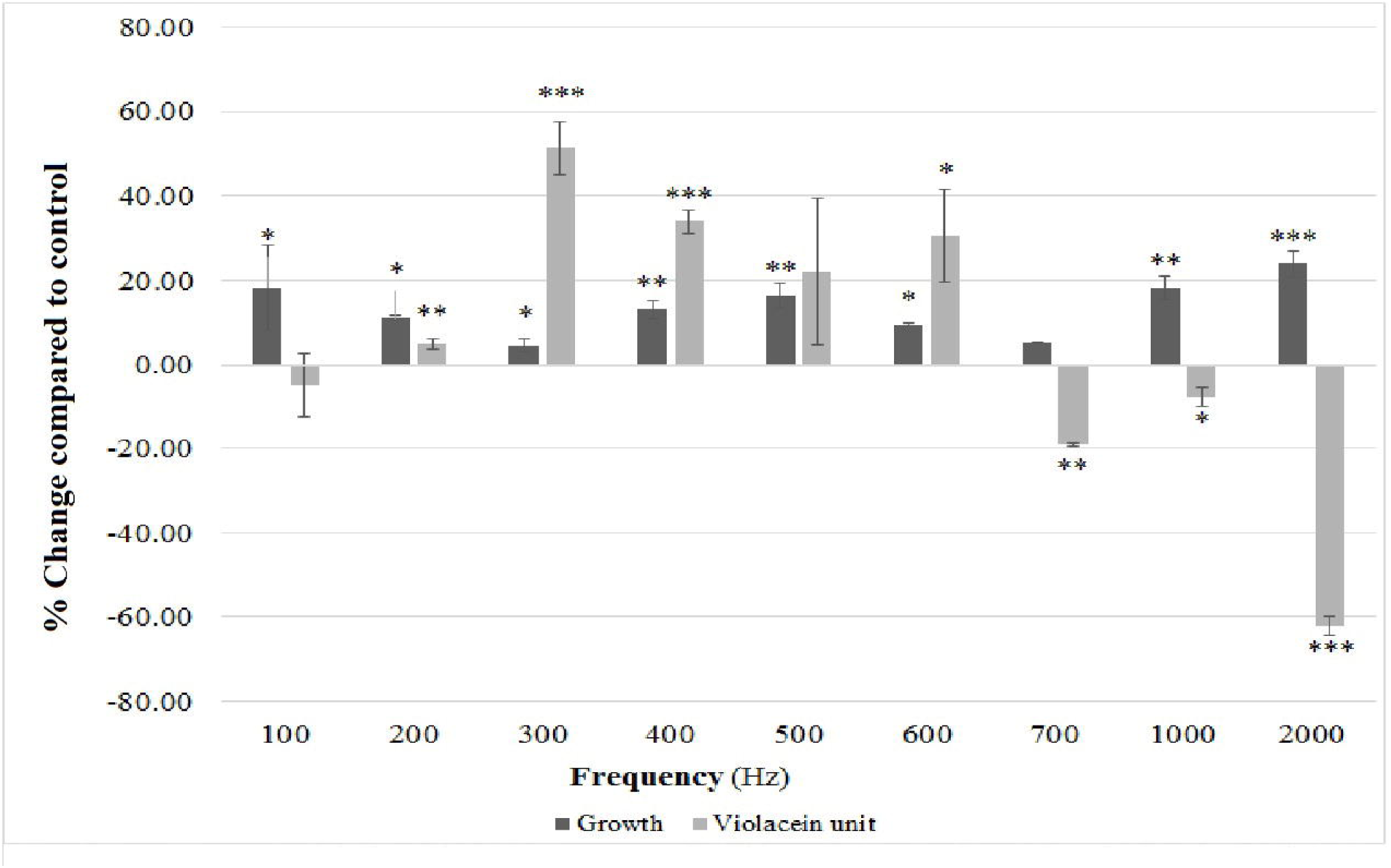
Effect of various sonic frequencies on growth and violacein production of *C. violaceum*. Bacterial growth was measured as OD_764;_ OD of violacein was measured at 585 nm, and violacein unit was calculated as the ratio OD_585_/OD_764_ (an indication of violacein production per unit of growth); \**p*<0.05, \*\**p*<0.01, \*\*\**p*<0.001; Control: Culture receiving no sonic stimulation, Experimental: Culture receiving sonic stimulation

In this study selection of the test frequency was based on our observation during our previous studies (Sarvaiya and Kothari, 2015; Shah et al., 2016), where we found that sound (corresponding roughly to a frequency range of 50-750 Hz) applied in form of music could notably alter growth and production of certain metabolites by the test microorganisms. 1000 and 2000 Hz frequencies were selected based on reports from other researchers. For example, Gu et al. (2016) reported 2,000 and 8,000 Hz sound to enhance biomass and growth rate of *Escherichia coli* significantly. 2000 Hz is believed to be one of the major frequency component in natural world (Cai, 2013).

### 3.2. Differential gene expression in C. violaceum exposed to 300 Hz sound

Whole transcriptional profiling of the *C. violaceum* culture resulted in identification of a total number of 7,403 coding regions. A total of 425 gene were significantly (p<0.05) differentially expressed, of which 342 were found to be up-regulated and 83 were found to be down-regulated. However, applying the dual criteria for screening i.e. p<0.05, and a cut-off value of 1.5 for fold change, none of the down-regulated genes (list provided as supplementary information) were found to be significant. All the up-regulated genes passed both the significance criteria. List of all the differentially up-regulated genes is provided in Table 2, including the 143 hypothetical proteins whose functions are not known. Function-wise categorization of the significantly up-regulated genes is presented in Figure 3. One of the top 20 up-regulated genes was a member of the LuxR family of transcriptional regulators. In total, 2 such LuxR family transcriptional regulators were found to be up-regulated. Violacein synthesis in the sound-stimulated culture was found to increase by 1.52 fold, and as violacein synthesis in this organism is under control of QS, we expected differential expression of the genes, which are part of the QS circuit of *C. violaceum*. Though up-regulation was observed for the LuxR analogue (i.e. CviR), no up-regulation was observed for the AHL synthase. This indicates that there may not be any enhanced signal production in sonic-stimulated culture, but more number of the AHL binding protein CviR are synthesized, leading to the formation of more number of CviR-AHL complex (not leaving much unused AHL), which in turn may bind to their target promoters, and cause overexpression of the relevant genes. Intriguingly, a hypothetical gene CV_2441 probably coding for dienelactone hydrolase was also upregulated, which is an AHL-degrading enzyme (Hong *et al.*, 2012), whose effect is likely to be quorum quenching. The gene CV_3413 coding for an acyl carrier protein was found to be 2.59 fold up-regulated. Acyl carrier protein has been considered as the key component of bacterial fatty acid synthesis. A key feature of the fatty acid synthetic pathway is that all of the intermediates are covalently bound to a protein called acyl carrier protein (ACP), a small, very acidic and extremely soluble protein. However, none of the genes involved in fatty acid synthesis were found to be up-regulated. It seems that a good amount of ACP flux was diverted in the sound stimulated culture towards polyketide synthesis, as the polyketide synthase (CV_4293) was also up-regulated. ACP, the central cofactor protein for fatty acid synthesis, supplies acyl chains for lipid A and lipoic acid synthesis, as well as quorum sensing, bioluminescence and toxin activation. Furthermore, ACPs or PCPs (peptidyl carrier proteins) are also utilized in polyketide and non-ribosomal peptide synthesis, which produce important secondary metabolites (Chan and Vogel, 2010).

**Figure 3.**
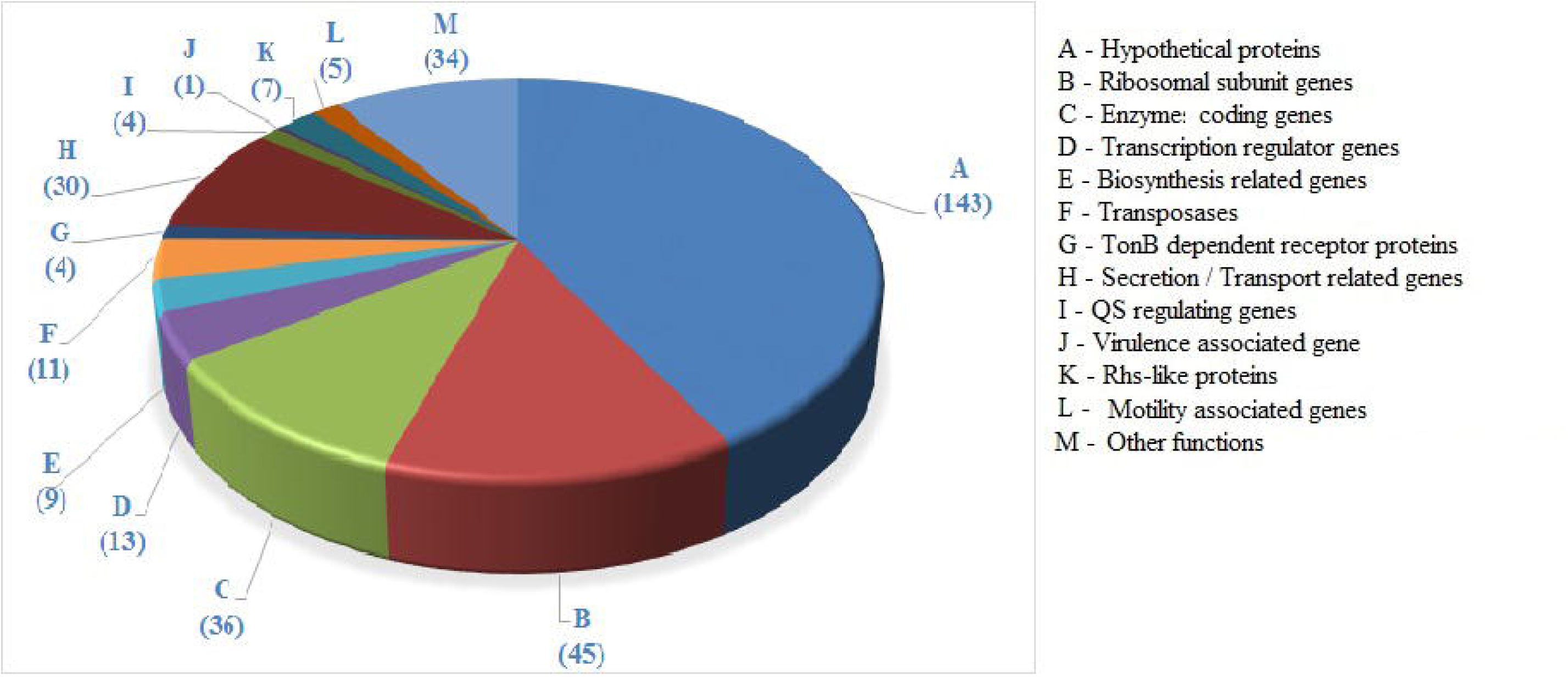
Function-wise categorization of the significantly up-regulated genes.

External sonic stimulation can be considered a sort of stress for the test bacterium, and under this situation the stress response machinery (e.g. the enzymes involved in ROS detoxification) of the organism can be expected to be up-regulated. One such stress adaptation protein disulfide oxidase CV_3998 (coded by *dsbA*) was found to be up-regulated in this study by 1.91 fold. Ability to cope with such environmental stress is believed to stem from the plethora of specific transporters present in this bacterium (Vasconcelos *et al.*, 2004), and we did find a good number (nearly 30) of proteins involved in transport and/or secretion to be significantly up-regulated. Many of them may be making it possible for the sonic-stressed cells to uptake nutrients at a faster pace, followed by their faster utilization. For example, a 3-fold up-regulated glucose 6-phosphate dehydrogenase (G6PD) (CV_0145), and 3.16-fold upregulation of nucleotide sugar epimerase can be taken as an indication of enhanced glucose metabolism through the pentose phosphate pathway. Interestingly, these enzyme activities are essential for later formation of erythrose-4-phosphate (E4P), which is the starting point of synthesis of the aromatic amino acids. One of these aromatic amino acids, tryptophan is the precursor for violacein synthesis (Antonio and Pasa, 2004). *C. violaceum* can convert a larger amount of glucose to aromatic amino acids than other organisms, owing to its lack of a complete hexose monophosphate pathway (HMP). HMP-defective mutants have been shown to produce an increased amount of E4P, which is the limiting substrate for tryptophan biosynthesis. Tryptophan hyper-producing *Corynebacterium glutamicum* were shown to possess a modified pentose phosphate pathway (Ikeda and Katsumata, 1999).

The reaction catalyzed by the overexpressed G6PD generates reducing power in form of NADPH. Biosynthesis of violacein requires NADPH for conversion of L-tryptophan in cell-free extracts; one or more of the NADPH binding sites in VioA, VioC, and VioB may be responsible for this requirement (Hoshino and Yamamoto, 1997; August et al., 2000). Requirement for NADPH has been demonstrated for at least two enzymes (VioD and VioC) participating in violacein biosynthesis (Xu et al., 2016), and thus increased availability of NADPH stemming from overexpressed G6PD can have a direct positive impact on violacein biosynthesis.

Among the genes of the *vio* operon, the one coding for polyketide synthase (i.e. *vio* B) was found to be up-regulated by 3.92 fold, which is a rate limiting enzyme of the violacein synthesis pathway (Antônio and Creczynski-Pasa, 2004; Xu et al., 2016). VioB is a large multifunctional enzyme, speculated to catalyze the 1,2-indole shift of tryptophan as well as the condensation reaction to generate the violacein-pyrrole ring (August *et al.*, 2000). Another enzyme, 5-aminolevulinate synthase (CV_0803), which was 2-fold up-regulated is involved in the pathway leading to the formation of the natural tetrapyrrole pigments.

In this study, cell yield was found to increase up to a minor extent only, whereas violacein production was enhanced at a much higher magnitude. This suggests that violacein production is not that tightly related to (or regulated by) cell density. Literature review reveals that QS signals may not be the sole control mechanism in *C. violaceum* for violacein production. Even in densely populated cell cultures, grown in high glucose and oxygen levels, violacein production can be suppressed (Antônio and Creczynski-Pasa, 2004). Violacein production thus, can also be considered to be dependent on the carbon source and its concentration in the culture medium, indicating possibility of a control mechanism similar to that mediated by cyclic AMP.

In total, four kinases (CV_0024, CV_3542, CV_4059, CV_1789) were found to be up-regulated indicating an enhanced phosphorylation. It may be noted here that protein kinases play an important role in regulation of enzyme activities within a cell (Berg et al., 2007). Multiple genes (including 9 *Sec* genes, and 2 Type III secretion proteins) associated with secretion were found to be up-regulated. This indicates a notable increase in the secretion activities of the sound stimulated culture. This fact may be looked in context of a large number (45) of up-regulated genes coding for ribosomal proteins, which can be taken as an indication of enhanced protein synthesis. Higher protein synthesis may cause the cell to secrete more protein content to the exterior.

Sound waves travel through the liquid medium containing bacteria as a mechanical vibration. These vibrations are likely to increase the oscillations among the microbial population, by promoting the probabilistic collision between the intracellular molecules, thereby increasing the magnitude of the systemic noise. Inside the cells, intracellular processes inevitably involve collisions between molecules such as DNA, mRNA, and proteins (Patnaik, 2012), and the sonic treatment is very much likely to enhance the impact of such collisions.

In total, five genes [*fliP, fliC, flag* (2), *cheY*] associated with chemotaxis/motility/flagella were found to be up-regulated in sound-stimulated culture, indicating an increased mobility of the *C. violaceum* cells. Chemotaxis is believed to help the survival of cells. One of the essential chemotaxis associated genes *cheY* (upregulated 2.54 fold) in its phosphorylated form diffuses through the cytoplasm and binds to the flagellar motor switch. The bound CheY~P functions as an allosteric regulator governing the rotations of the flagellar motor. Enhanced membrane permeability and faster glucose uptake in the sound–stimulated culture can be thought to result in fluctuations in the chemo-attractant concentrations, and interfere with the chemosensory response, finally affecting chemotaxis. Such fluctuations will represent a source of noise. This noise can affect intracellular processes having a significant effect on cell motility. As the Che proteins of the bacteria are expressed by specific genes, noise in gene expression can have a notable influence on the chemotactic motility. Noise associated with gene expression can be intrinsic or extrinsic. Cell-intrinsic noise can arise from fluctuations in cell-specific factors, viz. metabolite (e.g. E4P, violacein) concentrations. Internally driven fluctuations can be propagated from one cell to another. Faster nutrient uptake in the sound-stimulated culture (Shah et al., 2016) is likely to cause a rapid change in extracellular concentration of chemoattractants, making the bacteria more responsive to these changes. Intracellular noise in bacterial cell may be present at level of transcription or translation. Translational noise has been shown to have an influence on gene expression (Ozbudak et al., 2002), and this noise can be believed to be at a higher level in sound-stimulated culture owing to an increased protein synthesis. Sonic stimulated culture seems to be characterized by an increased chatter (rapid fluctuations in the rotation of the flagellar motors).

One of the major focus of this study was to see effect of sonic stimulation on QS of the bacterial population. QS is a mechanism to regulate population behaviour. The collective cellular response may be related to the impact of noise. The behavioral variability of a cell population can be controlled by the slope of a histidine kinase activation curve (Emonet and Cluzel, 2008). In the sound-stimulated culture, a total of four histidine kinase associated genes were found to be up-regulated. Of which two were two-component sensors coding for histidine kinase. Phosphorylation of a histidine residue occurs in response to detection of an extracellular signal, to initiate a change in cell state or activity (http://amigo.geneontology.org/amigo/term/G0:0000155). In context of the present study, this extracellular or environmental change can be considered to be the sound wave. The remaining two up-regulated genes are regulatory proteins involved in auto phosphorylation of histidine kinase. Of which, CheY is a chemotaxis regulator protein, and another is LuxR family transcriptional regulator.

Overall biosynthesis, metabolism and enzyme activities seem to be up-regulated in the sound-receiving culture. Higher metabolism and higher enzyme activity will increase the demand for energy, and hence these cells may be expected to produce and consume more ATP than control cells. This is in line with the observed up-regulation (1.97 fold) of ATP synthase subunit delta (*atpH*), and GTPaseObg (2.29 fold) involved in ATP synthesis (i.e. energy availability) and GTP hydrolysis (i.e. energy release), respectively. The most effective sound signal found in this study was 300 Hz. A nearby frequency of 340 Hz could drive ATP formation in bovine F_1_/F_0_ ATPase function in sub-mitochondrial particles (Syroeshkin *et al.*, 1998). Sound can be believed to be capable of altering ATP rotation or proton gradients across membrane, provided that other transmembrane enzymes and transporters are also susceptible to the sonic vibrations (Aggio et al., 2012). It should be noted here that ten (including 5 porins) transport associated genes were up-regulated in our experimental culture.

A liquid growth medium disturbed by travelling of the sound waves through it can be said to offer a high level of external (environmental) noise to the cells growing in it, and this external noise inevitably exerts an impact on the ligand binding dynamics of the bacterial population in question. Optimum levels of noise can generate more efficient chemotaxis, which seems to be the case here, as indicated by up-regulation of chemotaxis/motility related genes. Conventionally, systems performance is measured by signal-to-noise ratio. However, this ratio may not be a fitting metric for biological systems. It has been suggested that if the aim is to evaluate the effect of noise on one or more biological functions, it is more useful to measure variations in those very functions (McDonnell and Abbott, 2009), e.g. cell yield and violacein concentration measured in this study. This study has clearly demonstrated a notable degree of differential gene expression in sound treated bacterial culture. Fluctuations in the gene expression level, rates of metabolic reactions, and the concentrations of participating reactants and products can contribute to a rise in intrinsic noise (Kaern et al., 2005; Raser and O’shea, 2005).

Though effect of sonic vibration on yeast metabolome has been reported previously (Aggio et al., 2012), to the best of our awareness, this is the first report describing the altered gene expression in bacteria subjected to sonic range of sound waves. This study has demonstrated the capacity of audible sound signal to affect bacterial growth and QS-regulated pigment production. Differential up-regulation of a large number (342) of genes (4.63% of *C. violaceum* genome) was found to occur in the sound-stimulated culture. In the bacterial culture receiving sonic treatment, particularly the genes associated with protein synthesis, chemotaxis, motility, various enzyme activities, E4P synthesis, transcriptional regulators, and histidine kinase, were upregulated. The gene expression profile of this sound-stimulated culture can be said to be associated with violacein overproduction, and this information can be useful for strain improvement. Such studies providing clear demonstrations of the ability of sonic waves to affect microbial metabolism can open new perspectives at the interfaces of acoustics, microbiology, biophysics, and biochemistry. In long term, such studies can pave the way for development of sonic stimulation as a tool for manipulating microbial growth and metabolism.

## 4. Conclusions

Among the nine different sonic frequencies tested, 300 Hz sound enhanced violacein production by *C. violaceum* the most. A total of 342 genes were significantly up-regulated in the *C. violaceum* culture exposed to 300 Hz sound treatment. To the best of our awareness, this is the first report describing differential gene expression in bacteria exposed to sonic stimulation, and this information can be of direct use in strain improvement of *C. violaceum* for enhanced production of the bioactive metabolite violacein.

Table 1. Level of sound intensity at different frequency

Table 2. List of significantly up-regulated genes

## Acknowledgement

Authors thank GUJCOST (Gujarat Council on Science and Technology) for financial support, and Nirma Education and Research Foundation (NERF) for infrastructural support.

